# IsoformSwitchAnalyzeR: Analysis of changes in genome-wide patterns of alternative splicing and its functional consequences

**DOI:** 10.1101/399642

**Authors:** Kristoffer Vitting-Seerup, Albin Sandelin

## Abstract

Alternative splicing is an important mechanism involved in both health and disease. Recent work highlights the importance of investigating genome-wide changes in patters of splicing and the subsequent functional consequences. Unfortunately current computational methods only support such analysis on a gene-by-gene basis. To fill this gap, we extended IsoformSwitchAnalyzeR thereby enabling analysis of genome-wide changes in both specific types of alternative splicing as well as the predicted functional consequences of the resulting isoform switches. As a case study, we analyzed RNA-seq data from The Cancer Genome Atlas and found systematic changes in both alternative splicing and the consequences of the associated isoform switches.

**Availability:** Windows, Linux and Mac OS: http://bioconductor.org/packages/IsoformSwitchAnalyzeR.

**Contact:** KVS, AS

## 1 Introduction

Alternative splicing, alternative transcription start- and termination sites (here, for simplicity, jointly referred to as alternative splicing) expand the RNA repertoire of most human genes (Forrest *et al.*, 2014; The ENCODE Project Consortium, 2012). Changes in which isoform(s) that are used in different conditions are common, and examples of such isoform switches with important functional consequences have been described in many biological processes (reviewed in (Baralle and Giudice, 2017; Urbanski *et al.*, 2018)). This has motivated genome-wide screen for isoform switches with predicted functional consequences, resulting in identification of hundreds of genes (Sebestyen *et al.*, 2015; Vitting-Seerup and Sandelin, 2017; Climente-González *et al.*, 2017). The abundance of isoform switches may reflect important genome-wide changes and indeed large scale changes, such as systematic shortening of 3’UTRs in cancers, have previously been reported (e.g. (Miura *et al.*, 2013; Xia *et al.*, 2014)). Despite these findings most studies only focus individual genes and/or isoform switches. The lack of global analyses may reflect the lack of suitable computational tools; to our knowledge, there are no tools for genome-wide analysis of patterns of alternative splicing or the resulting isoform switches. To this end, we extended IsoformSwitchAnalyzeR, originally designed for exploration of individual isoform switches (Vitting-Seerup and Sandelin, 2017), with the ability to analyze changes in large-scale patterns of alternative splicing and isoform switch consequences.

## 2 Methods

The extensions to the easy-to-use R package IsoformSwitchAnalyzeR are available in version >1.1.10. Splicing classification is described in Supplementary Fig. 1 and (Vitting-Seerup *et al.*, 2014). Enrichment tests are performed via base R’s prop.test and comparisons of enrichments are done with fisher.test. *P* values are corrected for multiple testing using the Benjamin-Hochberg scheme and a *FDR* < 0.05 is considered significant. Additional details can be found in the documentation via Bioconductor. For the example below, Colon Adenocarcinoma (COAD) and Liver Hepatocellular Carcinoma (LIHC) RNA-Seq data from The Caner Genome Atlas (TCGA) (Chang *et al.*, 2013) was extracted from (Vitting-Seerup and Sandelin, 2017). Analysis downstream of the isoform switch identification was redone with IsoformSwitchAnalyzeR v 1.1.0 and only genes containing isoform switches with the predicted consequences shown in (Fig.1A) were considered. All data analyses and associated plots presented here are directly produced by IsoformSwitchAnalyzeR; no custom analysis or image post-processing was performed.

**Figure 1:**
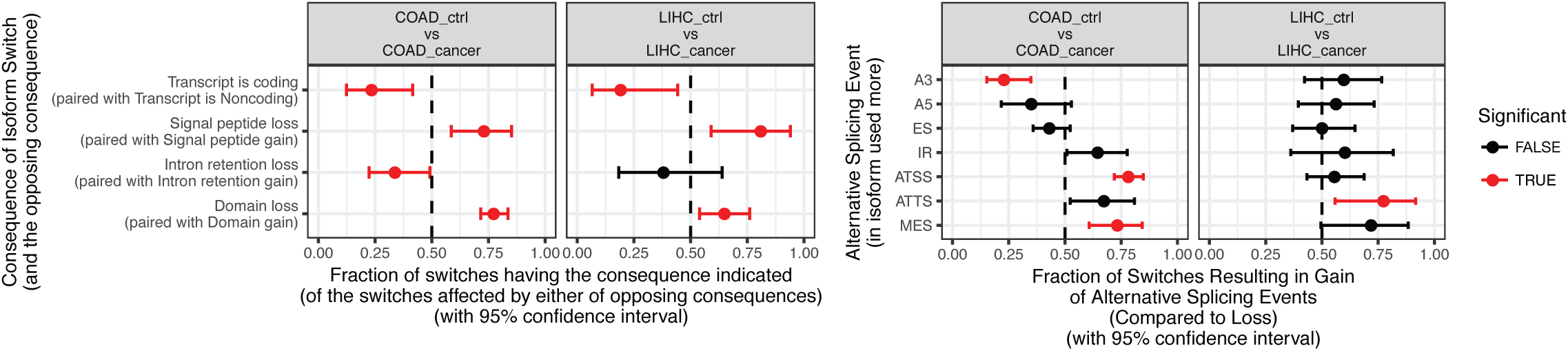
Enrichment/depletion in isoform switch consequences (A, by the extractConsequenceEnrichment function) and alternative splicing (B, by the extractSplicingEnrichment function) where subplots indicate each of two cancer types analyzed. **A**) From isoform switches resulting in either gain/loss of a consequence, the x-axis shows the fraction (with 95% confidence interval) resulting in the consequence indicated by y axis, in the switches from ctrl to cancer. **B**) The fraction (and 95% confidence inter-val) of isoform switches (x-axis) resulting in gain of a specific alternative splice event (indicated by y axis) in the switch from ctrl to cancer. Dashed line indicate no enrichment/depletion. Color indicate if FDR < 0.05 (red) or not (black). Splicing abbreviation are as in Figure S1. Note the figures are exactly as produced by IsoformSwitchAnalyzeR via the indicated functions.

## 3 Results

To showcase the identification and analysis of changes in genome-wide patterns of alternative splicing and isoform switch consequences we applied IsoformSwitchAnalyzeR to RNA-seq data from TCGA. Specifically, we analyzed COAD and LIHC by comparing each to their respective adjacent normal tissue controls (ctrl), resulting in the identification of hundreds of individual genes with isoform switch consequences and alternative splicing events (Supplementary Fig. 2 A-B).

To analyze large-scale patterns in predicted isoform switch consequences, we extracted all isoform switches resulting in a gain/loss of a specific consequence (e.g. protein domain gain/loss) when comparing cancer and ctrl. We then tested for enrichment or depletion by testing whether the gain/loss ratio was different from the expected 1:1, using a standard proportion test. This analysis, and a visual representation of the results, was all done using the extractConsequenceEnrichment() function in IsoformSwitchAnalyzeR. We found that many types of isoform switch consequences were either enriched or depleted in isoform switches between respectively normal controls and COAD and LIHC cancer samples (Fig. 1A). For example, compared to normal, loss of protein domains occurred much more frequently than expected in both cancer types (Fig. 1A, all *FDR* < 0.05, proportion test). Intriguingly, COAD and LIHC had highly similar patterns of enrichment/depletions of isoform switch consequence types (Fig. 1A). To assess this observation, we used the extractConsequenceEnrichmentComparison function, which performs a statistical and a visual comparison of the enrichment analysis in the individual cancer types (which was described above). We could verify the similarity of the enrichment/depletion patterns of the two cancer types as no significant differences were found (Supplementary Fig. 2 C, all *FDR* >= 0.1, Fishers exact test).

Analysis of genome wide-changes in alternative splicing can be performed using the same approach as above. This is done by analyzing the fraction isoform switches resulting in the gain of a specific alternative splicing event (e.g. exon skipping gain) an analysis also implemented in IsoformSwitchAnalyzeR. To illustrate this, we used the extractSplicingEnrichment function to analyze COAD and LICH for enrichment/depletion of alternative splicing events. Given the similarity of the pattern of isoform switch consequence enrichment/depletion in the two cancer types (Fig. 1A) we were surprised to observe that the enrichment/depletion patterns of splicing events were substantially different (Fig. 1B). We used the extractSplicingEnrichmentComparison() function to compare the enrichments of alternative splicing patterns in COAD and LICH and found that the differences in enrichment of alternative 3’ acceptor site and alternative transcription start sites were statistically significant between the two cancer types (Supplementary Fig. 2 D, FDR < 0.001, Fishers exact test). Thus, the observed isoform switch consequences, although similar between cancers, may result from different underlying biological mechanisms.

## 4 Conclusion

IsoformSwitchAnalyzeR enables analysis of changes in genome-wide patterns of alternative splicing and isoform switch consequences. We illustrated this functionality by analysis of TCGA data, and found that certain switch consequence and splice patterns are specifically enriched/depleted.

## Funding

This work has been supported by grants from the Lundbeck Foundation and the Independent Research Fund Denmark

## Conflict of Interest

None.

---

**Figre S1.**
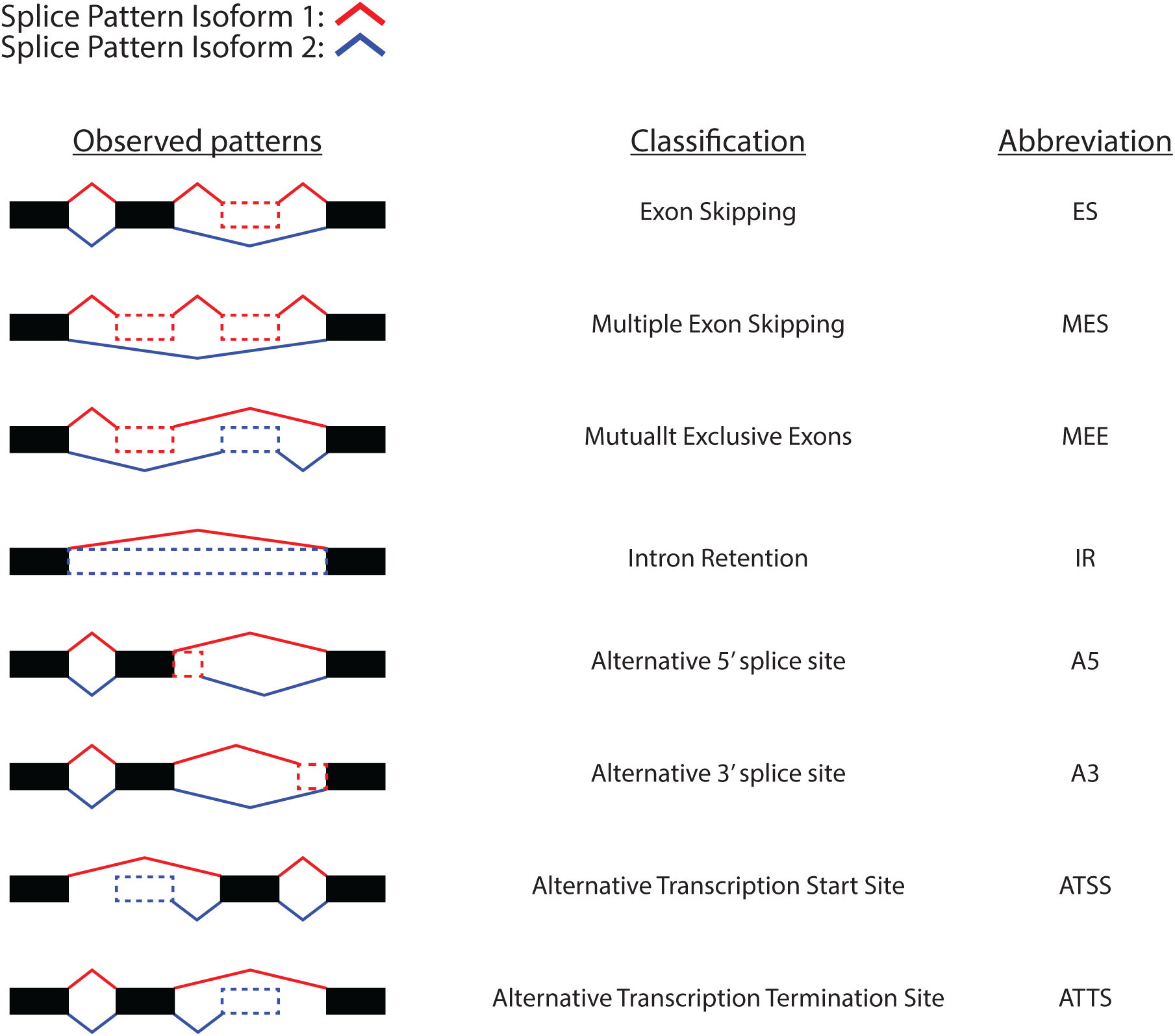
The observed splice patterns (left colum) of two isoforms compared as indicated by the color of the splice patterns. The corresponding classification of the event (middle column) and the abreviation used (right column). Inspired by (Vitting-Seerup et al., 2014)

**Figure S2:**
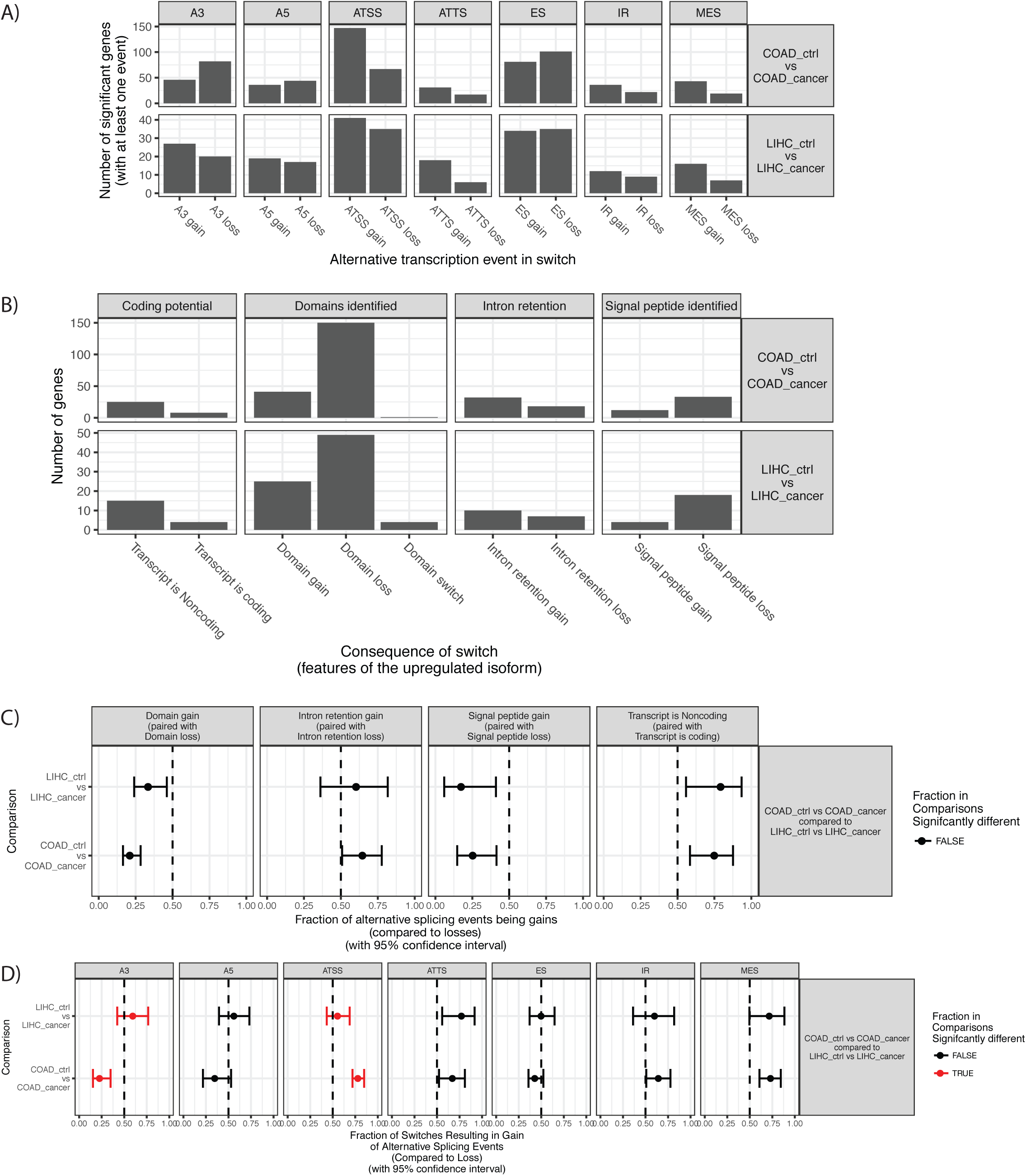
**A, B**: Number of isoforms significantly differentially used between cancer and ctrl (rows of sub-plots) resulting in at least one splice event (A) or isoform switch consequence (B) (colums of subplots). A is produced by the extractSplicingSummary function. B is produced by the extractConsequenceSummary function. **C, D**: Comparison of enrichment/depletion in isoform switch consequences in the two cancer types (y-axis) (C, by the extract-ConsequenceEnrichmentComparison function) and alternative splicing (D, by the extractSplicingEnrichmentComparison function) where subplots indicate consequence (C) or alternative splicint (D) type. C) From isoform switches resulting in either gain/loss of a consequence, the x-axis shows the fraction (with 95% confidence interval) resulting in the consequence indicated by y axis, in the switches from ctrl to cancer. D) The fraction (and 95% confidence inter-val) of isoform switches (x-axis) resulting in gain of a specific alternative splice event (indicated by y axis) in the switch from ctrl to cancer. Dashed line indicate no enrichment/depletion. Color indicate if FDR < 0.05 (red) or not (black). Note the figures are exactly as produced by the IsoformSwitchAnalyzeR via the indicated functions.

